# PCC1 treatment reshapes ribosomal, lysosomal and membrane-lipid transcriptional programs in therapy-induced senescent human stromal cells

**DOI:** 10.64898/2026.08.17.745159

**Authors:** Qun Wang, Jie Li, Qinghua Lyu

## Abstract

Cellular senescence combines stable proliferative arrest with extensive changes in secretory, metabolic and organelle programs. Procyanidin C1 (PCC1) has dose-dependent senomorphic and senolytic activity, but the structure of its transcriptome-wide response is not well defined. We reanalyzed published RNA-sequencing counts from bleomycin-induced senescent PSC27 human stromal cells treated with 50 µM PCC1. The primary contrast contained three independent biological replicates per condition. PCC1 altered 8,420 genes at adjusted *P*<0.05 and absolute log2 fold change≥1. Ribosomal genes showed the clearest coordinated response, followed by lysosomal genes. Analysis across the full ontologies also identified lysosomal-membrane, proton-transport, late-endosome, lipid-localization, cholesterol and fatty-acid programs. Prespecified analyses revealed two trajectories relative to senescence. Lipid-transport, lipid-binding and plasma-membrane programs decreased during senescence and increased after PCC1, whereas lysosomal-membrane, mitochondrial-membrane, cholesterol and ion-transport programs increased in both contrasts. Senescence-associated outputs were more selective. A prespecified SASP-effector score decreased, but a broad Reactome SASP set did not pass false-discovery correction. AP-1-family expression shifted, and an HSP90/HSF1/proteostasis panel increased. Together, these data support a model in which PCC1 induces coordinated transcriptomic remodeling in senescent stromal cells, encompassing membrane-lipid remodeling, changes in cellular infrastructure, and selective modulation of senescence-associated outputs.

## Introduction

Cellular senescence is a stress-responsive cell state characterized by durable proliferative arrest and broad remodeling of chromatin, metabolism, organelles and intercellular signaling [1]. A prominent but non-obligate feature is the senescence-associated secretory phenotype (SASP), which can contain cytokines, chemokines, growth factors, proteases and extracellular-matrix regulators [2,3]. SASP composition varies with the initiating insult, cell lineage, time and tissue environment. This heterogeneity enables senescent cells to support repair and immune surveillance in some settings while sustaining chronic inflammation or tumor-promoting niches in others [1–3]. Senescence is therefore better understood as a family of related states than as a single transcriptional endpoint.

Several regulatory and organelle systems help maintain the senescent state. AP-1 factors shape parts of the senescence enhancer landscape and contribute to a transcriptional program that retains substantial plasticity [4]. Mitochondrial dysfunction can produce a distinct secretory phenotype [5], while declining proteostasis reduces the ability of senescent cells to manage proteotoxic and oxidative stress [6]. Lysosomes are also integral to this state. Senescence-associated β-galactosidase reflects expansion of lysosomal mass and increased lysosomal GLB1 activity [7,8]. Spatial coupling of mTOR, autolysosomes and protein synthesis can sustain high secretory output [9], and the lysosomal proteome contributes selected components to the SASP [10]. Lysosomal membrane damage can create metabolic dependencies that are exploitable for senolysis [11]. These findings connect biosynthesis, mitochondrial state, lysosomal processing and secretory output in senescent cells.

PCC1 is an epicatechin trimer identified in a natural-product screen for senotherapeutic activity. In the original PSC27 study, lower concentrations attenuated SASP expression, whereas higher concentrations preferentially reduced senescent-cell viability through reactive-oxygen and mitochondrial responses [12]. Subsequent studies reported PCC1 activity in aged retina, pulmonary fibrosis, kidney fibrosis and aged hematopoietic or immune compartments [13–16]. Activity was also reported in skin fibrosis and vascular senescence [17,18]. Proposed mechanisms differ among models and include PUMA–BAX, ANGPTL4–NOX4, CEBPB, EGFR–TGF-β/SMAD and HSP90–TLR2/NF-κB signaling. PCC1 has also been linked to matrix-producing cancer-associated fibroblasts and to microbiome–FOXO1 regulation of the intestinal mucosal barrier [19,20]. Earlier studies in macrophage and neuronal systems reported context-dependent modulation of TLR4–MAPK/NF-κB and Nrf2–HO-1 signaling [21,22]. No single pathway is therefore likely to explain PCC1 activity across all experimental settings.

Here, we examined the PCC1 response at five connected levels: global transcription, ribosomal programs, lysosomal programs, membrane-lipid biology and selected senescence-associated outputs. We used the complete treatment contrast and tested whether the available control-group structure affected the estimates. Differential expression was integrated with ranked GO, Reactome and deposited-author KEGG annotations. We also examined prespecified membrane-lipid axes in the primary contrast and in an exploratory senescence contrast. SASP and proteostasis panels were evaluated alongside broader gene sets and direct AP-1-family expression. This analysis identified coordinated ribosomal, lysosomal and membrane-lipid changes together with a more selective response among senescence-associated outputs.

## Results

### PCC1 produces a broad transcriptional shift that is stable across models

We reanalyzed count data from the published PSC27 cell experiment and RNA-sequencing study [12]. In the source experiment, primary human prostate stromal cells were exposed to bleomycin, allowed to develop therapy-induced senescence and then treated with 50 µM PCC1. We performed no cell culture or RNA sequencing; the present study reanalyzes the deposited counts (Fig. 1a). The complete primary contrast included independent biological replicates of bleomycin-induced senescent cells (BLEO) and PCC1-treated senescent cells (BLEO+PCC1).

**Figure 1.**
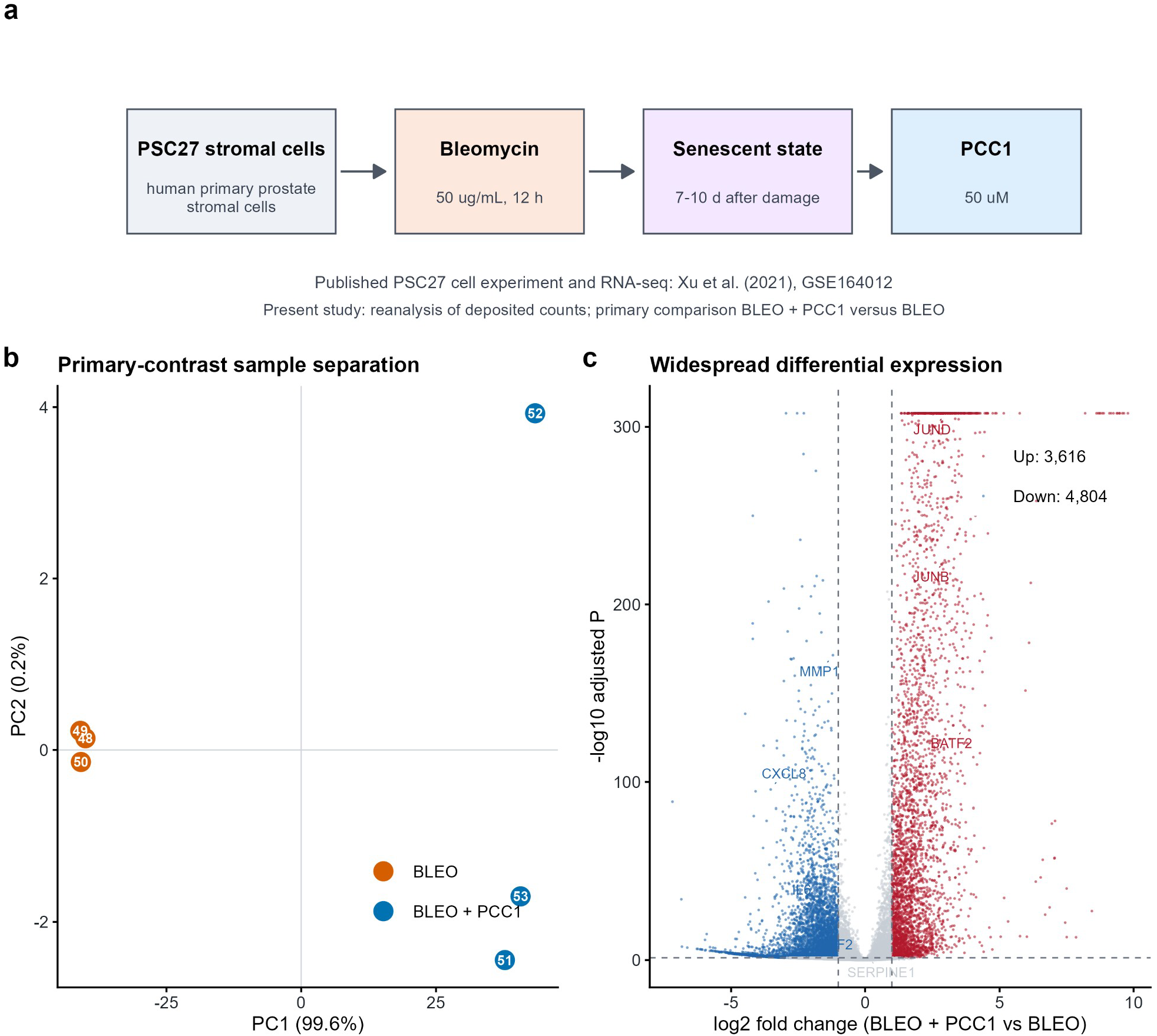
Published experimental design and the PCC1-associated global transcriptomic transition. **a**, Source experiment and analysis design. Xu and colleagues performed the PSC27 cell experiment and RNA sequencing [12]: primary human prostate stromal cells were exposed to bleomycin (50 µg mL^-1^ for 12 h), maintained for 7–10 days to establish senescence and treated with 50 µM PCC1. The present study performed no cell experiment or sequencing; it reanalyzed deposited GSE164012 counts, with BLEO+PCC1 versus BLEO as the primary contrast. **b**, Principal-component analysis of variance-stabilized expression for the six primary samples (n=3 independent biological replicates per condition). PC1 and PC2 explain 99.6% and 0.2% of variance, respectively. **c**, Volcano plot for BLEO+PCC1 versus BLEO. Increased and decreased genes meet adjusted *P*<0.05 and absolute log2 fold change≥1. Extremely small adjusted *P* values were visually capped on the y axis; uncapped values remain in the source table. The analysis identified 3,616 increased, 4,804 decreased and 9,032 other genes.

After filtering, 17,452 genes entered the differential-expression model. Principal-component analysis of variance-stabilized expression separated the two treatment groups almost entirely along PC1, which accounted for 99.6% of the variance; PC2 accounted for 0.2% (Fig. 1b). Samples remained tightly grouped within each condition. At the adjusted-*P* and fold-change thresholds, 3,616 genes increased and 4,804 decreased after PCC1 exposure. Another 9,032 genes did not meet both thresholds (Fig. 1c). The response was broad and bidirectional rather than confined to inflammatory markers.

The deposited design expected three untreated controls, whereas the locally available NCBI combined count matrix contained two. We designated the complete BLEO versus BLEO+PCC1 contrast as primary and used an eight-sample model containing the available controls only for sensitivity analysis. Effect estimates changed negligibly between the models. Spearman ρ was 0.999991 for log2 fold changes and 0.999152 for Wald statistics. Effect direction agreed for 99.90% of genes (Supplementary Fig. S1). Inclusion of the incomplete untreated-control group therefore had little effect on the central PCC1 contrast.

### Ribosomal programs form a dominant and internally consistent PCC1-associated signal

Ranked enrichment identified a broad cellular-infrastructure response, with translation and ribosome terms among the strongest signals (Fig. 2 and Supplementary Fig. S2). GO Cellular Component analysis reproduced this pattern in the six-sample primary contrast. Large ribosomal subunit (NES=2.035, FDR=1.08×10^-8^) and ribosomal subunit (NES=2.008, FDR=1.08×10^-8^) were strongly enriched. Cytosolic large ribosomal subunit, cytosolic ribosome and ribosome had NES values of 2.005, 1.954 and 1.906, respectively. Each had FDR=1.08×10^-8^ after PCC1 (Fig. 2a). Mitochondrial ribosome was also positively enriched (NES=1.782, FDR=1.53×10^-5^), showing that the response extended beyond cytosolic ribosomes.

**Figure 2.**
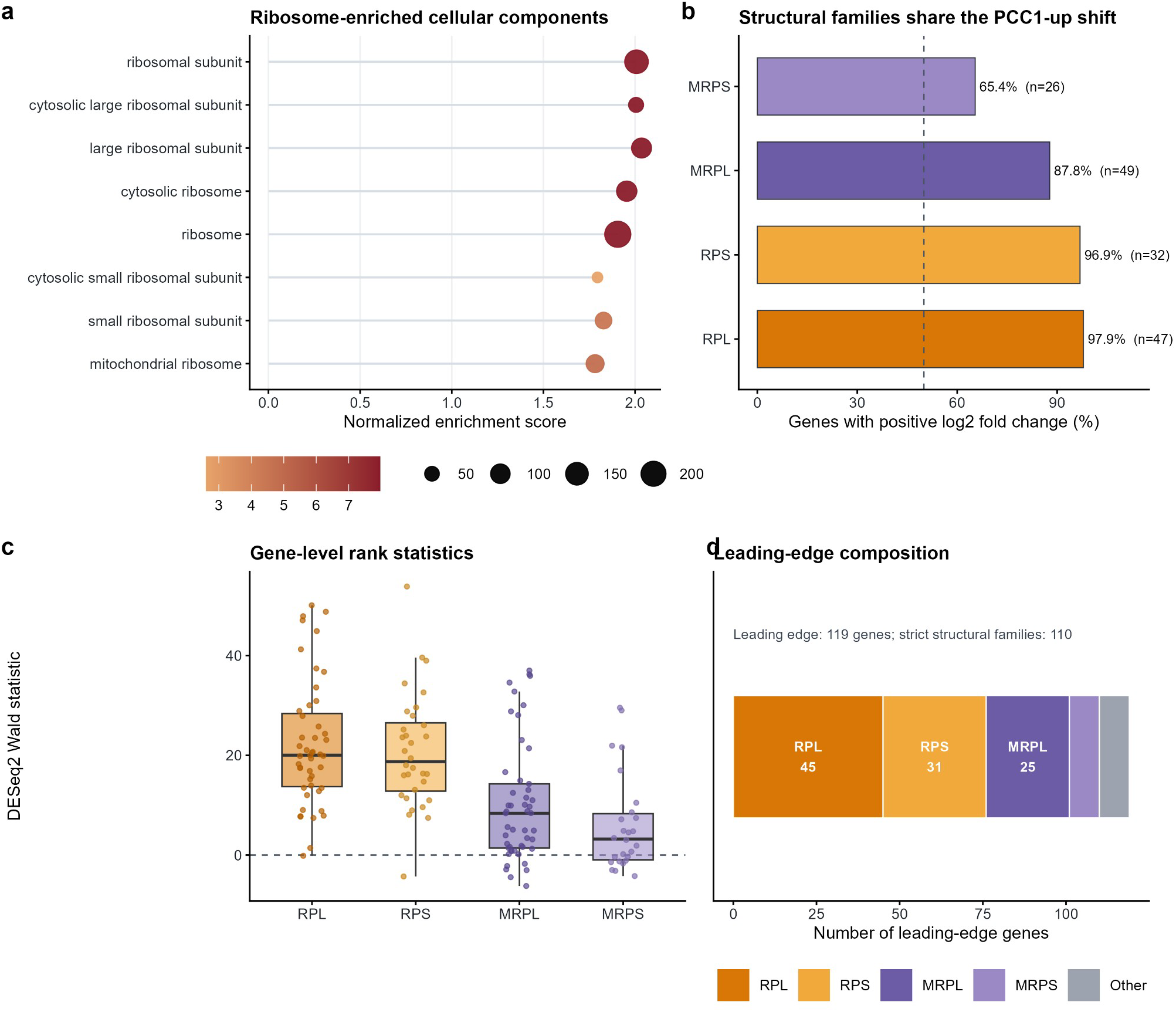
Ribosomal remodeling is broad, directional and robust across structural families. **a**, Selected GO Cellular Component ribosomal terms from six-sample preranked enrichment. Position denotes NES, point size denotes gene-set size and color denotes -log10 FDR. **b**, Percentage of strict structural ribosomal-family genes with positive log2 fold change. The dashed line marks 50%; percentages and quantified family sizes are shown. **c**, Gene-level DESeq2 Wald-statistic distributions for RPL, RPS, MRPL and MRPS families. Boxes show medians and interquartile ranges; points are individual genes. **d**, Composition of the 119-gene leading edge for the ribosomal-subunit term. Strict structural families account for 110 genes. Family definitions used protein-coding symbols matching RPL, RPS, MRPL or MRPS patterns. Complete values are provided in the panel source tables.

Gene-family analysis showed that this enrichment was not driven by a small subset of unusually responsive transcripts. Among strict structural families, 97.9% of RPL genes and 96.9% of RPS genes had positive log2 fold changes. The corresponding proportions were 87.8% for MRPL and 65.4% for MRPS genes (Fig. 2b). The distributions of gene-level Wald statistics were shifted upward across all four families, with the largest median shifts in the cytosolic RPL and RPS groups (Fig. 2c). Of 119 genes in the leading edge of the ribosomal-subunit enrichment, 110 belonged to strict structural ribosomal families, comprising 45 RPL, 31 RPS, 25 MRPL and 9 MRPS genes (Fig. 2d). Gene-set enrichment, family-level direction, gene-level ranks and leading-edge composition all supported the same ribosomal response.

The ribosomal response occurred within a wider infrastructure program. Cytoplasmic translation was enriched in GO Biological Process (NES=1.707, FDR=6.96×10^-7^). The deposited-author KEGG ribosome set showed NES=2.129 and FDR=3.10×10^-8^. Oxidative phosphorylation was enriched in GO (NES=1.784, FDR=3.48×10^-6^) and in the deposited-author KEGG collection (NES=1.823, FDR=3.20×10^-7^). Reactome terms for translation elongation, ribosome-associated quality control, mitochondrial translation, respiratory electron transport and mitochondrial protein import changed in the same positive direction (Supplementary Fig. S2). Agreement across annotation systems linked ribosomal remodeling to concurrent biosynthetic and bioenergetic changes.

### Lysosomal, vacuolar and endosomal programs define a second major axis of remodeling

Lysosome-associated enrichment was detected in four annotation contexts (Fig. 3a). The deposited-author KEGG lysosome pathway was positively enriched (NES=1.825, FDR=6.09×10^-7^). GO Biological Process identified lysosomal lumen acidification (NES=1.775, FDR=0.0173) and vacuolar acidification (NES=1.688, FDR=0.0206). GO Cellular Component identified lysosomal membrane, lytic vacuole membrane, vacuolar lumen and lysosomal lumen. All had positive NES values, and the membrane and lumen terms had FDR below 10^-6^. Reactome late endosomal microautophagy was also positively enriched (NES=1.658, FDR=0.0279). The agreement among pathway, process and compartment annotations supports broad lysosomal remodeling.

**Figure 3.**
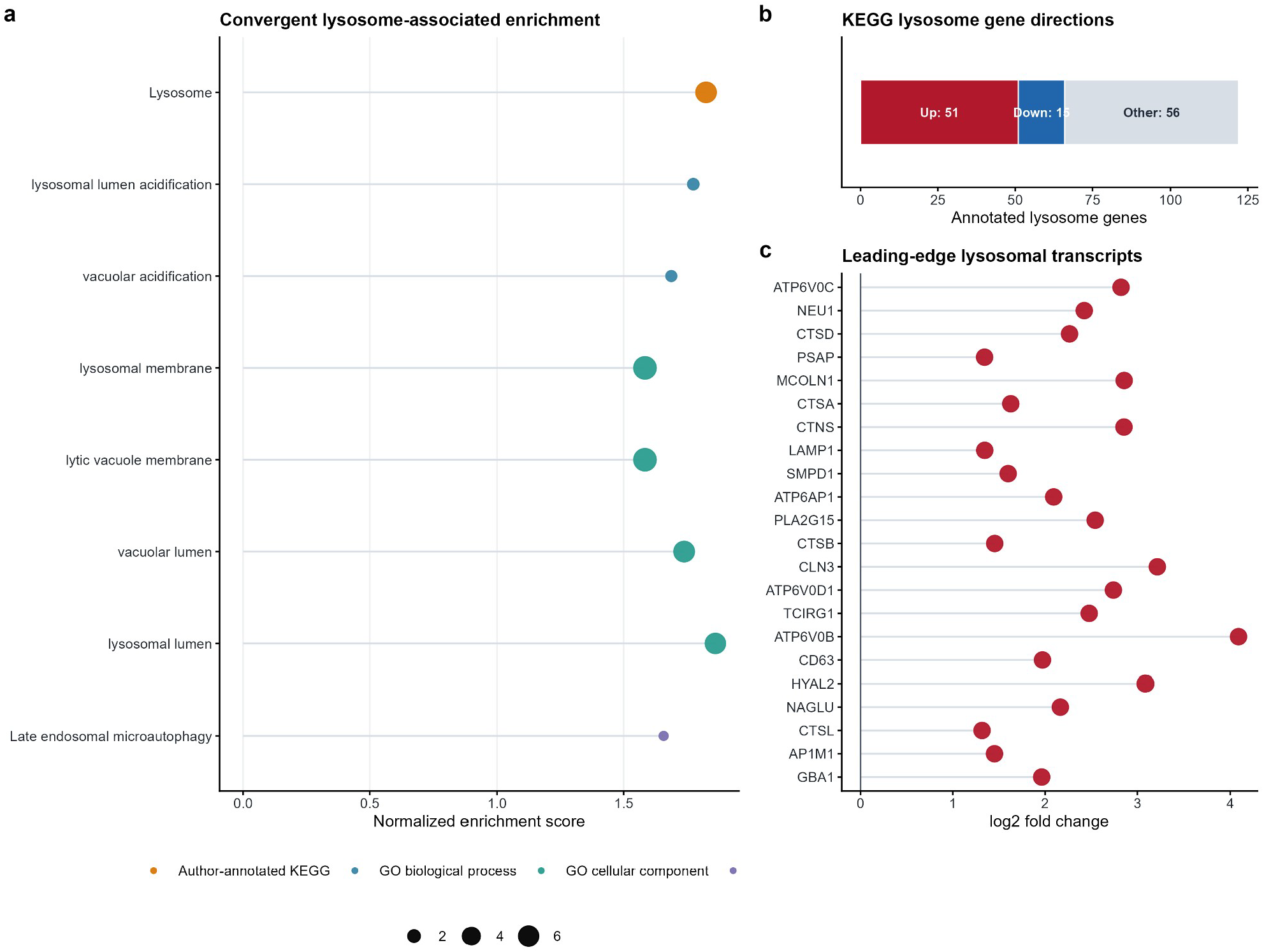
PCC1 is associated with coordinated lysosomal, vacuolar and endosomal transcriptional remodeling. **a**, Significant positive lysosome-, vacuole- or endosome-associated terms from deposited-author KEGG, GO Biological Process, GO Cellular Component and Reactome analyses. Position denotes NES; color identifies the annotation source and point size denotes -log10 FDR. **b**, Differential-expression status of 122 quantified genes in the deposited-author KEGG lysosome set. Up and Down require adjusted *P*<0.05 and absolute log2 fold change≥1; Other genes do not meet both thresholds. **c**, Top significant transcripts in the positive KEGG lysosome leading edge, ranked by absolute Wald statistic. Position denotes log2 fold change and point size denotes -log10 adjusted *P*. Genes include acidification machinery, membrane and trafficking components, and lysosomal hydrolases. Panel c displays contributors to enrichment rather than an independently tested gene panel.

The annotated KEGG lysosome set contained 122 quantified genes. Fifty-one met the increased-expression thresholds, 15 met the decreased-expression thresholds and 56 did not meet both adjusted-*P* and fold-change cutoffs (Fig. 3b). The positive leading edge included multiple functional classes required for lysosomal capacity. V-ATPase and acidification components included ATP6V0B, ATP6V0C, ATP6V0D1, ATP6AP1 and TCIRG1. Membrane and trafficking genes included LAMP1, CD63, CTNS, CLN3, MCOLN1 and AP1M1. Hydrolases or accessory proteins included CTSA, CTSB, CTSD, CTSL, GBA1, NAGLU, NEU1, PSAP, SMPD1, HYAL2 and PLA2G15 (Fig. 3c). Several of these transcripts increased by more than two log2 units, and ATP6V0B increased by 4.09 log2 units.

A single enrichment score does not capture the lysosomal response. It includes lumen and membrane components, proton-pump machinery, trafficking factors and catabolic enzymes. Together with the increased ribosomal and respiratory programs, this pattern suggests coordinated changes in protein production, energy supply and degradative capacity. The transcriptomic data cannot distinguish increased flux, compensatory lysosomal biogenesis or preferential survival of cells with greater lysosomal capacity. Each explanation makes a different experimental prediction.

### Membrane and lipid-state remodeling separates restorative from amplified trajectories

The primary full-ontology analysis extended the lysosomal signal into membrane and lipid biology (Fig. 4a). Lysosomal membrane was strongly enriched after PCC1 (NES=1.588, FDR=1.08×10^-8^). Proton transmembrane transporter activity (NES=1.671, FDR=2.72×10^-4^) and late endosome membrane (NES=1.458, FDR=0.00932) were also enriched. Lipid localization, unsaturated-fatty-acid metabolism, lipid droplets, sterol metabolism, cholesterol metabolism and fatty-acid metabolism all passed FDR<0.05. FDR was calculated within each complete relevant GO ontology, not within a membrane-only subset.

**Figure 4.**
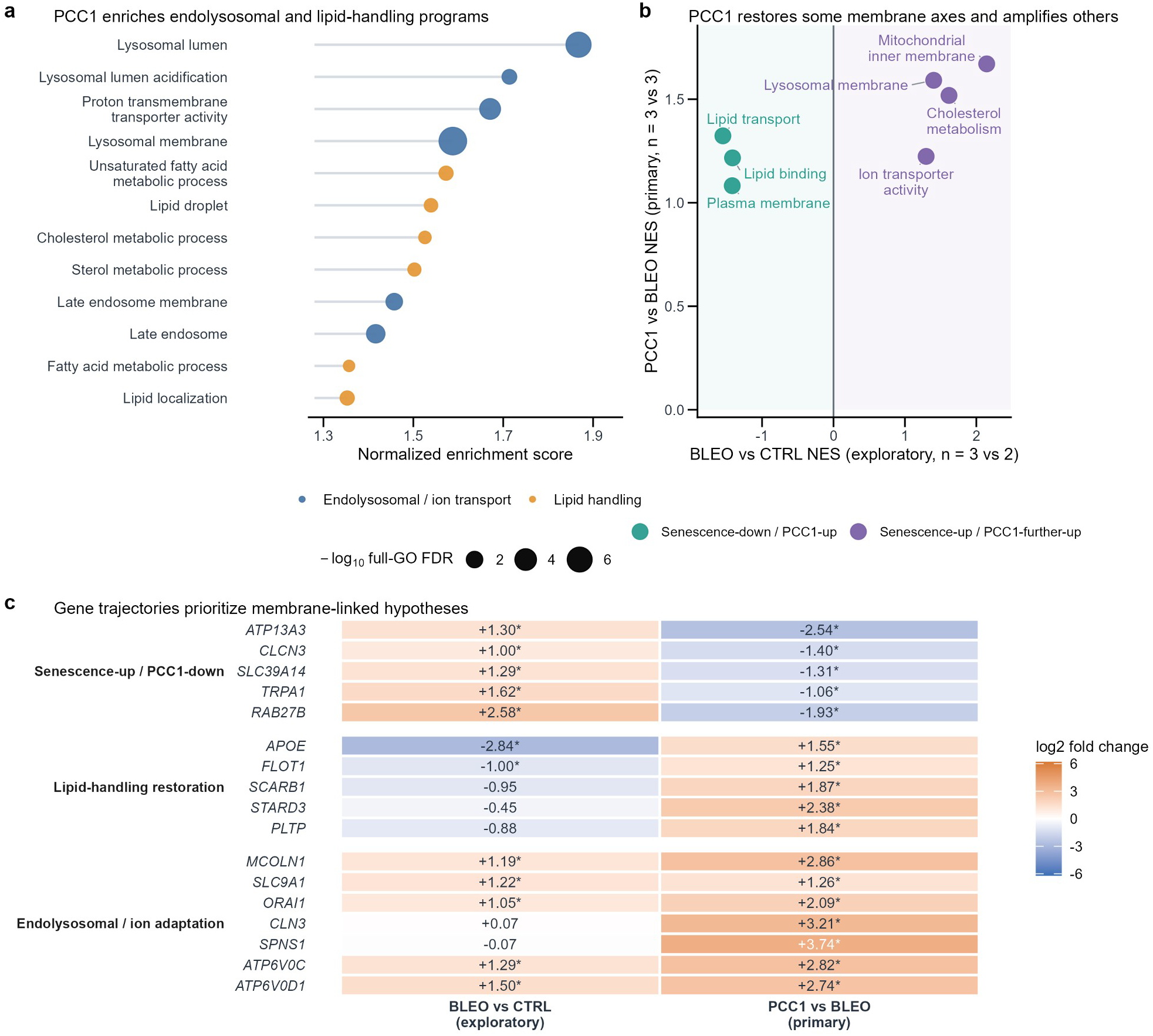
PCC1 remodels membrane-lipid programs through restorative and amplified trajectories. **a**, Selected membrane- and lipid-related terms from full GO Biological Process, Molecular Function and Cellular Component ranked enrichment for primary PCC1 versus BLEO (n=3 independent biological replicates per condition). Position denotes NES, point size denotes -log10 FDR and color denotes biological class. FDR was calculated within each complete GO ontology. **b**, Predefined-axis trajectories for exploratory BLEO versus CTRL (n=3 versus n=2) and primary PCC1 versus BLEO (n=3 versus n=3). Teal axes decreased during senescence and increased after PCC1. Purple axes increased in both contrasts. All displayed axes passed FDR<0.05 in both comparisons, with correction across the predefined axes within each contrast. **c**, Selected candidate-gene log2 fold changes. Asterisks denote adjusted *P*<0.05 and absolute log2 fold change≥1 within the relevant genome-wide differential-expression analysis. Candidate selection was post hoc and is hypothesis-generating. Panel b is exploratory because the deposited count matrix contains only two CTRL samples. The figure supports membrane-lipid state remodeling, not direct binding or a low-dose mechanism.

Directional over-representation of strict differentially expressed genes provided an independent summary of the primary contrast. Increased genes were over-represented in lysosomal membrane (odds ratio=2.02, FDR=3.80×10^-9^) and lipid metabolic process (odds ratio=1.56, FDR=2.78×10^-8^). Lipid transport (odds ratio=1.70, FDR=1.59×10^-4^) and cholesterol metabolism (odds ratio=2.36, FDR=2.58×10^-4^) were also enriched. These FDR values were calculated across 25 predefined axes in both DEG directions.

An exploratory BLEO-versus-CTRL comparison then separated two senescence-relative trajectories (Fig. 4b). Lipid transport, lipid binding and plasma-membrane programs decreased in BLEO cells and increased after PCC1. By contrast, lysosomal membrane, mitochondrial inner membrane, cholesterol metabolism and ion-transporter activity increased in both comparisons. The control group contained two samples, so this trajectory analysis is hypothesis-generating rather than confirmatory.

Gene-level trajectories highlighted candidates for experimental follow-up without identifying direct PCC1 targets (Fig. 4c). ATP13A3, CLCN3, SLC39A14, TRPA1 and RAB27B increased during senescence and decreased after PCC1. APOE and FLOT1 showed the opposite pattern. MCOLN1, SLC9A1, ORAI1, ATP6V0C and ATP6V0D1 increased in both comparisons, while CLN3 and SPNS1 increased strongly after PCC1. These candidates link lipid handling, ion homeostasis, endolysosomal adaptation and vesicle export to the observed transcriptional state.

### PCC1 selectively remodels SASP, AP-1 and proteostasis programs

We next asked whether the global response produced uniform suppression of senescence-associated transcription. A prespecified 15-gene SASP-effector panel comprised SERPINE1, MMP12, CSF2, CXCL2, IL1A, CXCL3, CCL20, CXCL1, CCL2, IL6, PTGS2, IL1B, MMP3, CXCL8 and MMP1. The sample-level mean decreased from 10.550 VST units in BLEO cells to 9.499 after PCC1. The difference was -1.051 VST units (exploratory Welch *P*=8.72×10^-5^; Fig. 5a). This coordinated decrease supports attenuation of a compact group of inflammatory and matrix-remodeling effectors.

**Figure 5.**
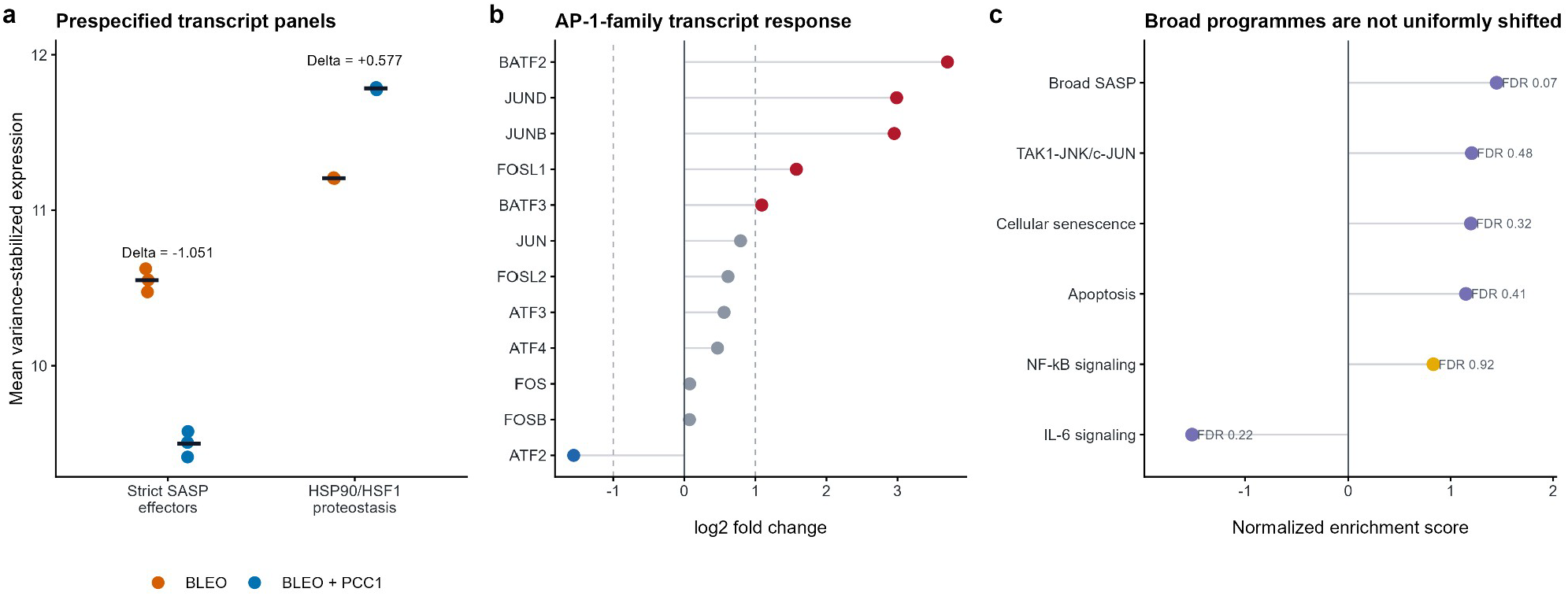
PCC1 selectively remodels SASP, proteostasis and AP-1-associated transcription. **a**, Per-sample variance-stabilized scores for a prespecified 15-gene SASP-effector panel and a 15-gene HSP90/HSF1/proteostasis panel (n=3 independent biological replicates per condition). Points are biological replicates; horizontal bars indicate means. Delta denotes BLEO+PCC1 minus BLEO. Welch *P* values are exploratory nominal tests. **b**, AP-1-family log2 fold changes from the primary DESeq2 contrast. Red and blue points meet adjusted *P*<0.05 and absolute log2 fold change≥1; gray points do not meet both thresholds. **c**, Ranked enrichment of broader senescence-associated and mechanistic programs. Text labels give FDR values; none passes FDR<0.05, including Reactome SASP (FDR=0.071). The panels distinguish a reduced canonical effector module from heterogeneous broad senescence and AP-1-family responses.

Broader annotations did not simply extend the compact-panel result. The Reactome SASP set had positive ranked enrichment (NES=1.449, nominal *P*=0.00544) but did not pass false-discovery correction (FDR=0.0710). Interleukin-6 signaling was negatively enriched but also non-significant after correction (NES=-1.517, FDR=0.225), and Reactome cellular senescence had NES=1.198 and FDR=0.317 (Fig. 5c). PCC1 reduced a defined SASP-effector core, while other genes in broad senescence annotations remained stable or shifted upward. The data therefore support selective output remodeling rather than broad suppression of the senescent transcriptome.

AP-1-family transcripts showed a similarly structured response (Fig. 5b). JUNB (log2 fold change=2.95), JUND (2.99), FOSL1 (1.58), BATF2 (3.70) and BATF3 (1.09) increased significantly, whereas ATF2 decreased (-1.56). JUN, FOSL2, ATF3 and ATF4 had smaller positive changes, while FOS and FOSB were not significant. Consistent with this mixed gene-level pattern, the Reactome program describing TAK1-mediated activation of JNK and c-JUN did not meet the FDR threshold (NES=1.206, FDR=0.476; Fig. 5c). The data identify a change in AP-1-family composition rather than uniform activation or suppression.

A prespecified 15-gene HSP90/HSF1/proteostasis panel increased by 0.577 VST units after PCC1 (exploratory Welch *P*=1.57×10^-5^; Fig. 5a). By contrast, broad apoptosis and inflammatory labels did not organize the ranked response at this time point. Reactome apoptosis had NES=1.149 and FDR=0.406; the deposited-author KEGG apoptosis and NF-κB pathways had FDR values of 0.232 and 0.916, respectively (Fig. 5c). Chaperone and stress-adaptive transcription therefore changed alongside ribosomal, mitochondrial and lysosomal programs. Apoptosis and NF-κB were not the dominant genome-wide responses in the measured cell population.

## Discussion

The PCC1-associated transcriptome separated into two broad components (Fig. 6). One comprised ribosomal, oxidative-phosphorylation, lysosomal and membrane-lipid programs. The other comprised a reduced SASP-effector core, altered AP-1-family composition and increased proteostasis transcripts. Broad SASP, senescence, apoptosis and NF-κB annotations were not uniformly suppressed. The magnitude and consistency of the first component indicate that cellular infrastructure is a major part of the response to PCC1 exposure.

**Figure 6.**
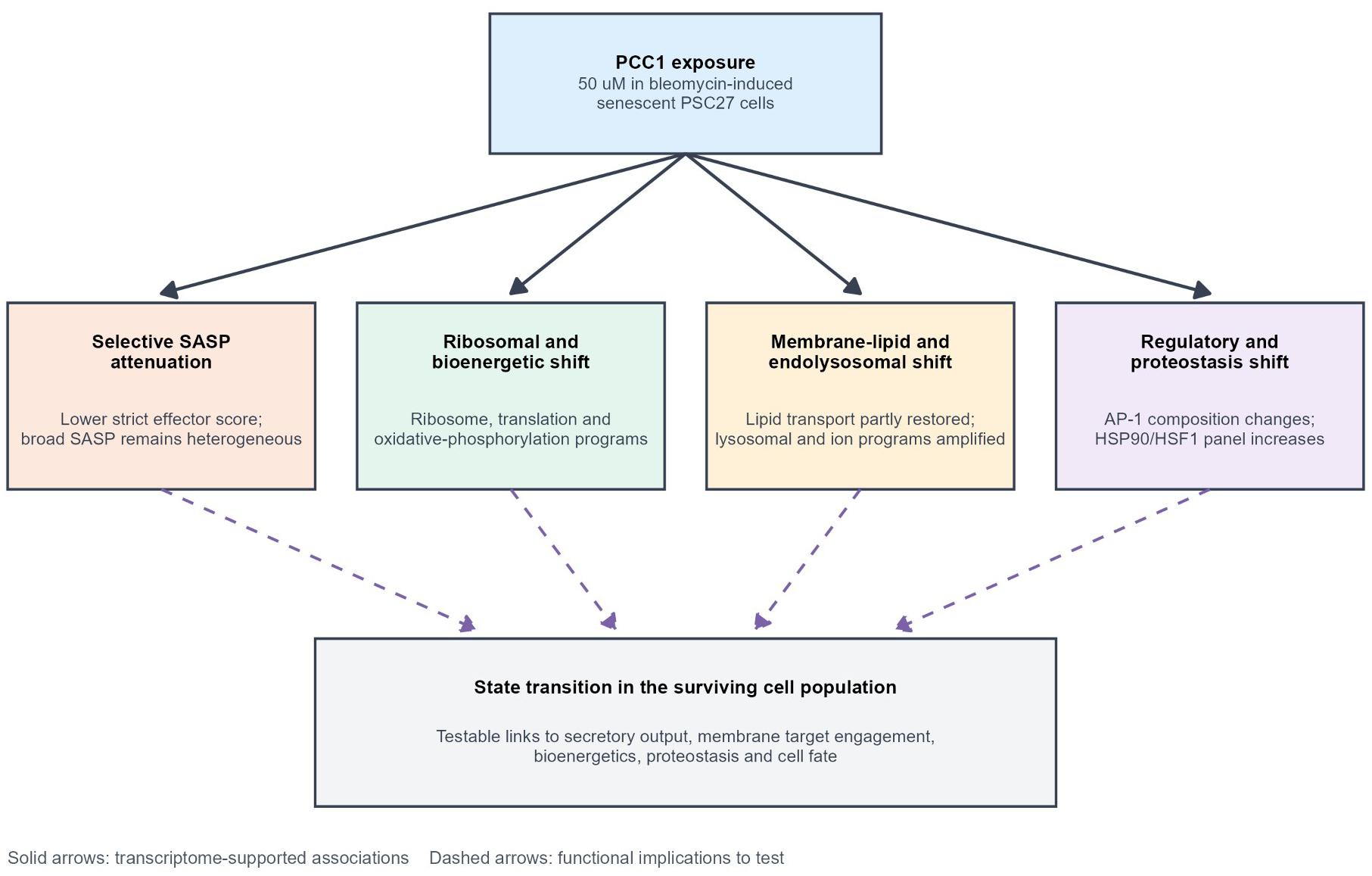
Evidence-layered model of the PCC1-associated transcriptional state. Solid connections summarize transcriptome-supported associations: strict SASP-effector attenuation; ribosomal and oxidative-phosphorylation remodeling; membrane-lipid and endolysosomal remodeling; AP-1-family reconfiguration; and increased proteostasis transcripts. Dashed connections denote functional implications to test. The model proposes that these modules jointly shape the surviving cell population. It does not assign causality, direct PCC1 binding or senescence-selective target engagement from enrichment alone.

Several independent observations support the ribosomal result. Positive enrichment recurred across GO, Reactome and deposited-author KEGG collections, almost every cytosolic RPL/RPS gene shifted upward, and the leading edge contained both cytosolic and mitochondrial structural families. The pattern indicates a coordinated change in transcripts that support ribosome production and translation. It may reflect stress adaptation, altered biosynthetic demand or the composition of cells remaining after treatment. These explanations differ, but all predict that translational capacity influences the PCC1 response. Nascent-protein synthesis, polysome profiling and time-resolved ribosome biogenesis measurements would distinguish increased translation from transcriptional compensation.

The lysosomal response is relevant because lysosomes contribute directly to senescent-cell biology. Increased lysosomal mass and GLB1 activity contribute to the classical SA-β-gal phenotype [7,8]. Autolysosome–mTOR spatial coupling can support high secretory protein synthesis [9], and lysosomal exocytosis contributes selected proteins to the senescence secretome [10]. Lysosomal membrane damage also lowers intracellular pH and creates a glutaminolysis-dependent survival mechanism in senescent cells [11]. In this context, the coordinated increase in V-ATPase subunits, membrane proteins, trafficking factors and hydrolases suggests that PCC1 affects a system already involved in senescent-cell maintenance. Lysosomal pH, mass, proteolytic capacity, autophagic flux, membrane integrity and exocytosis now require direct measurement; enrichment alone cannot determine how these functions changed.

Together, the ribosomal and lysosomal responses suggest possible coordination between protein production and disposal. Senescent cells can sustain substantial biosynthetic and secretory activity, and the TASCC provides a precedent for spatially coupling protein synthesis with autophagic degradation [9]. PCC1-associated increases in translation, respiration and lysosomal machinery may reflect an adaptive attempt to maintain proteome balance under treatment stress. Alternatively, PCC1 may select for surviving cells with high infrastructure capacity. A short time course before major viability changes, followed by paired measurements of translation, respiration and lysosomal flux, could distinguish induction from selection. Either result would identify cellular systems that influence senomorphic or senolytic responses to PCC1.

The membrane-lipid analysis narrows this infrastructure model. Plasma-membrane, lipid-transport and lipid-binding programs showed a senescence-down and PCC1-up trajectory. That pattern is consistent with partial state restoration. Lysosomal-membrane, mitochondrial-membrane, cholesterol and ion-transport programs instead increased in both comparisons. This second pattern is consistent with persistent stress adaptation or compensatory remodeling. Their coexistence suggests that 50 µM PCC1 captures a mixed transition rather than a simple reversal of senescence.

Independent chemical and biophysical studies make a membrane-proximal mechanism plausible, but they do not establish it in senescent PSC27 cells. Purified PCC1 inhibited isolated Na+/K+-ATPase with an IC50 of 4.5±0.8 µM, while docking proposed rather than proved binding sites [23]. All-atom simulations placed pure PCC1 at model plasma and mitochondrial membrane interfaces and predicted local cholesterol exclusion [24]. Related procyanidin dimers and trimers altered liposome surface properties and fluidity at low micromolar concentrations [25]. A separate liposome study supported interaction with phospholipid headgroups without showing senescence specificity [26]. DARTS, CETSA and microscale thermophoresis also supported EGFR binding in an aging-related skin-fibrosis model [17]. These independent observations justify testing membrane or membrane-protein engagement directly.

The transcriptomic data identify downstream changes that are consistent with this prior evidence. They do not show that PCC1 binds a senescence-specific lipid or membrane protein. They also cannot explain low-dose senomorphic activity because the dataset contains one 50 µM condition and one post-treatment time point. Direct target-engagement assays, fraction-resolved lipidomics and matched dose-time experiments are needed to establish the proposed causal order. The most discriminating design would compare proliferating and senescent PSC27 cells before viability changes occur.

The SASP findings further qualify the interpretation of PCC1. SASP composition varies across senescence triggers and cell types [2,3], so a compact effector panel and a broad pathway annotation need not move together. The reduction of the 15-gene panel agrees with the senomorphic effects reported in the source PSC27 study and in retinal and immune models [12,13,16]. The positive but FDR-borderline direction of the broad Reactome SASP set also shows that “SASP suppression” is not a binary description. PCC1 appears to reduce an inflammatory and matrix-remodeling core while preserving or increasing other senescence-associated transcripts. This selectivity could alter paracrine output without silencing the entire senescent program.

AP-1-family reconfiguration offers a plausible regulatory bridge between these layers. AP-1 factors organize senescence-associated enhancer architecture, but the family acts through combinatorial dimers with distinct binding and cofactor preferences [4]. Strong increases in JUNB, JUND, FOSL1 and BATF2, together with reduced ATF2 and stable FOS/FOSB, could redistribute AP-1-dependent transcription toward a different enhancer state. This model does not require AP-1 to be switched globally on or off. It predicts that AP-1 motif accessibility and occupancy will change at selected SASP, lysosomal and infrastructure genes. Time-resolved protein abundance, phosphorylation, chromatin accessibility and CUT&RUN or ChIP-seq for the induced subunits would test that prediction directly.

Proteostasis may provide another link between the observed programs. Senescent cells show diminished stress-responsive proteostasis [6], making chaperone systems important determinants of survival under additional stress. In vascular senescence, PCC1 was reported to bind HSP90 and inhibit an HSP90–TLR2/NF-κB axis [18]. The increased HSP90/HSF1-associated transcript panel in PSC27 cells is consistent with compensatory heat-shock signaling after chaperone perturbation. This possibility can be tested through PCC1 target-engagement assays, HSP90 ATPase measurements, HSF1 nuclear localization and client-protein stability in the same stromal model. The result links an independently reported target to a specific adaptive program, without assuming that the vascular mechanism also operates in PSC27 cells.

PCC1 responses vary markedly across published models. PCC1-associated apoptosis has been prominent in senescent pulmonary myofibroblasts and renal tubular epithelial cells [14,15], whereas aged retinal and immune tissues display combined senolytic and senomorphic responses [13,16]. Skin fibrosis has been linked to EGFR–TGF-β/SMAD signaling [17], and vascular senescence to HSP90–TLR2/NF-κB [18]. Other studies linked PCC1 to matrix-producing tumor fibroblasts [19] or microbiome-derived valeric acid and FOXO1 [20]. Non-senescent macrophage and neuronal models further show that PCC1 can modulate TLR4–MAPK/NF-κB or Nrf2–HO-1 signaling [21,22]. The absence of dominant apoptosis or NF-κB enrichment here is therefore compatible with those reports. It identifies the programs that predominate at this dose and time point in the surviving stromal-cell population.

This study is limited to one therapy-induced senescence model, a single transcriptomic time point and a complete treatment contrast with multiple independent biological replicates. The primary comparison was stable to model specification, although one expected untreated-control sample is absent from the combined NCBI count matrix. Ranked enrichment and panel scores identify coordinated transcriptional programs and generate testable hypotheses. Time-resolved perturbation and functional assays are needed to determine which responses are induced within cells and which reflect population selection.

PCC1 treatment was associated with broad changes in ribosomal, lysosomal and membrane-lipid programs in bleomycin-induced senescent human stromal cells. Membrane-linked responses included partial restoration of lipid-handling functions and further increases in endolysosomal or ion-homeostatic programs. These changes accompanied selective attenuation of canonical SASP effectors, altered AP-1-family expression and increased proteostasis transcripts. The data make membrane target engagement a plausible hypothesis, but do not establish it. Direct target-engagement, lipidomic and dose-time experiments are required to move from transcriptomic association to a causal mechanism.

## Methods

### Dataset provenance and experimental design

RNA-sequencing data were obtained from Gene Expression Omnibus accession GSE164012, linked to SRA project SRP299687 and the PCC1 study by Xu and colleagues [12]. PSC27 is a primary normal human prostate-derived stromal-cell population composed predominantly of fibroblastic cells. In the published source protocol, cells were exposed to bleomycin (50 µg mL^-1^ for 12 h), washed and maintained for 7–10 days to establish therapy-induced senescence. The transcriptomic PCC1 condition used 50 µM PCC1. The expected design comprised untreated control, bleomycin-induced senescent and PCC1-treated bleomycin-induced senescent cells, each with three biological replicates. The present study performed no cell culture, treatment or sequencing; it reanalyzed the deposited count data.

An Editorial Expression of Concern was issued for the source article in 2026 regarding a mouse image in Figure 6b and unavailable underlying data requested during editorial follow-up [27]. The notice does not identify GSE164012 or the deposited RNA-sequencing count matrix. We therefore report the notice transparently and restrict the present evidential claims to the independently reanalyzed files available from GEO/NCBI.

### Count matrix and sample inclusion

The NCBI GRCh38.p13 raw-count matrix and gene annotation table were used, following established principles for count-based RNA-sequencing analysis [28]. The combined matrix contained GSM4994846– GSM4994853 and lacked the expected untreated-control sample GSM4994845. All six samples in the primary contrast were present: GSM4994848–GSM4994850 (BLEO) and GSM4994851–GSM4994853 (BLEO+PCC1).

Genes were collapsed by Entrez Gene identifier, and integer counts were retained. Genes with at least 10 counts in at least three primary samples were included in differential-expression analysis.

### Differential expression and sensitivity analysis

Differential expression was performed in R with DESeq2 using a design of ∼ condition and the contrast BLEO+PCC1 versus BLEO [29]. Benjamini–Hochberg-adjusted *P* values controlled multiplicity. Genes were classified as increased or decreased when adjusted *P*<0.05 and log2 fold change was at least 1 or at most -1, respectively. Variance-stabilizing transformation with blind=FALSE was used for sample-level visualization, signature scoring and principal-component analysis.

A sensitivity DESeq2 model included the two locally available CTRL samples together with the six primary samples. Log2 fold changes and Wald statistics were compared between models by Spearman correlation, and effect-direction agreement was calculated across shared genes.

### Ranked gene-set enrichment

Genes were ranked by the DESeq2 Wald statistic, with positive values representing higher expression after PCC1. Ranked enrichment followed the GSEA framework [30] and was implemented with clusterProfiler-compatible workflows [31]. GO Biological Process and Cellular Component gene sets used Gene Ontology resources [32,33]. Reactome pathways used the Reactome knowledgebase [34]. KEGG-like pathways were tested with the per-gene pathway annotations in the deposited 2020 author workbook, with KEGG nomenclature traced to the established database framework [35]. Gene sets were restricted to 10–500 members. *P* values were adjusted within each collection by the Benjamini–Hochberg method. Deposited-author KEGG results are labelled explicitly because pathway membership reflects the annotation bundled with GSE164012 rather than current online KEGG curation.

The six-sample GO Cellular Component analysis used gseGO with ont=“CC”, Entrez identifiers, the DESeq2 Wald-statistic ranking and eps=0. The selected ribosomal terms were predefined by their GO descriptions. Strict structural ribosomal families were defined from protein-coding symbols matching RPL, RPS, MRPL or MRPS family patterns. Positive-direction proportions were calculated from the sign of the primary log2 fold change. Leading-edge genes were obtained from the ribosomal-subunit GO term.

### Lysosomal gene analysis

Lysosome-associated terms were selected from significant positive GO Cellular Component, GO Biological Process, Reactome and deposited-author KEGG results using lysosome-, vacuole- or endosome-related descriptions. The KEGG lysosome gene set was obtained from the deposited author annotation. Gene status was summarized using the genome-wide thresholds applied to differential expression. Leading-edge genes were obtained from the positively enriched KEGG lysosome result, joined to the primary DESeq2 table and ranked by the absolute Wald statistic for display. This analysis describes the transcripts contributing to enrichment and does not use the gene-status counts as an additional hypothesis test.

### Membrane and lipid-state analysis

The primary DESeq2 Wald statistic was tested against GO Biological Process, Molecular Function and Cellular Component gene sets. Gene sets contained 10–500 members, and Benjamini–Hochberg correction was applied within each complete ontology. Membrane- and lipid-related results were selected by predefined biological concepts, then reduced to non-redundant interpretable terms for display.

Twenty-five predefined axes represented membranes, lipid metabolism or transport, ion transport, receptors, endocytosis and vesicle biology. Directional over-representation used strict increased and decreased genes against the 17,452-gene tested universe. Benjamini–Hochberg correction was applied across 50 axis-direction tests. Ranked enrichment of these axes was also performed separately for BLEO+PCC1 versus BLEO and BLEO versus CTRL. FDR was controlled across the predefined axes within each contrast.

The BLEO-versus-CTRL analysis contained three BLEO and two CTRL samples because GSM4994845 was absent from the NCBI combined count matrix. It was used only to prioritize senescence-relative trajectories. Gene-level candidates were selected post hoc from the predefined axes using effect direction, adjusted *P* value, abundance and functional interpretability. Candidate display was hypothesis-generating and did not constitute an independent statistical test.

### Signature scores and AP-1-family analysis

The strict SASP-effector panel comprised SERPINE1, MMP12, CSF2, CXCL2, IL1A, CXCL3, CCL20, CXCL1, CCL2, IL6, PTGS2, IL1B, MMP3, CXCL8 and MMP1. The HSP90/HSF1/proteostasis panel comprised HSPB1, HSF1, DNAJB1, CDC37, STIP1, HSPA1A, HSP90AB1, BAG1, AHSA1, BAG3, HSPA1B, DNAJA1, HSP90AA1, HSPH1 and DNAJB4. Counts were filtered at 10 or more in at least two samples, variance-stabilized with blind=FALSE, and averaged across detected panel members for each sample. BLEO and BLEO+PCC1 means were compared with two-sided Welch tests. These panel-level *P* values are exploratory nominal statistics for two prespecified biological summaries.

AP-1-family genes were extracted directly from the primary DESeq2 table. Gene-level adjusted *P* values are those from the genome-wide analysis. Subunit transcript abundance was not converted into an AP-1 activity score.

### Statistical analysis and reproducibility

Biological replicate was the unit of analysis. Unless stated otherwise, enrichment significance was defined as FDR<0.05 and differential-expression significance as adjusted *P*<0.05 with absolute log2 fold change≥1. FDR families were defined separately for each full annotation collection or predefined-axis analysis. The exploratory BLEO-versus-CTRL contrast was not treated as equivalent to the complete primary contrast. Exact sample numbers, effect sizes and correction status are given in the Methods, source-data tables and figure legends. Figure generation used scripted R workflows. Each figure is accompanied by panel-level source tables and vector or high-resolution raster exports.

### Data and code availability

The RNA-sequencing dataset is publicly available under GEO accession GSE164012 and SRA project SRP299687. The publication package contains the differential-expression table, ranked gene list, complete GO Cellular Component results, panel-level source data, ESI tables, figure-generation code and an environment manifest. No unpublished wet-laboratory data are claimed.

## Author contributions

Conceptualization, Q.L.; investigation and data acquisition, Q.W. and J.L.; original draft preparation, Q.W. and J.L.; review and editing, Q.W., J.L. and Q.L. All authors read and approved the final manuscript and accept accountability for the work.

## Declaration of generative AI and AI-assisted technologies

During preparation of this manuscript, the authors used OpenAI ChatGPT and Codex (accessed August 2026) under author supervision to assist with English-language editing, manuscript organization, the development of analysis code and the preparation of plotting code. Statistical analyses were executed on the cited public dataset using the documented workflows. All figures were generated deterministically from the underlying numerical data; no generative-AI image synthesis or AI-based alteration of experimental images was used. The authors reviewed and validated all AI-assisted text, code, numerical outputs, citations, figures and interpretations, made all final scientific decisions, and take full responsibility for the content of the manuscript.

