## Supplemental Tables and Figures for "PCC1 treatment reshapes ribosomal, lysosomal and membrane-lipid transcriptional programs in therapy-induced senescent human stromal cells": PCC1_bioRxiv_Supplementary_Information.pdf

### File summary

This file contains Supplementary Figures S1 and S2 with legends, covering sample-level quality control, primary-versus-sensitivity model stability and cross-annotation pathway enrichment supporting the PCC1 transcriptomic reanalysis.

### Contents

Supplementary Figure S1 | Sample-level quality control and stability of the primary PCC1 contrast ...................... 2
Supplementary Figure S2 | Cross-annotation overview of cellular-infrastructure pathway enrichment .................. 3

### Associated data file

Supplementary Data S1 is supplied separately as PCC1_Supplementary_Data_S1.xlsx and contains sample metadata, full processed results, figure-level source tables, signature membership and model-concordance data.

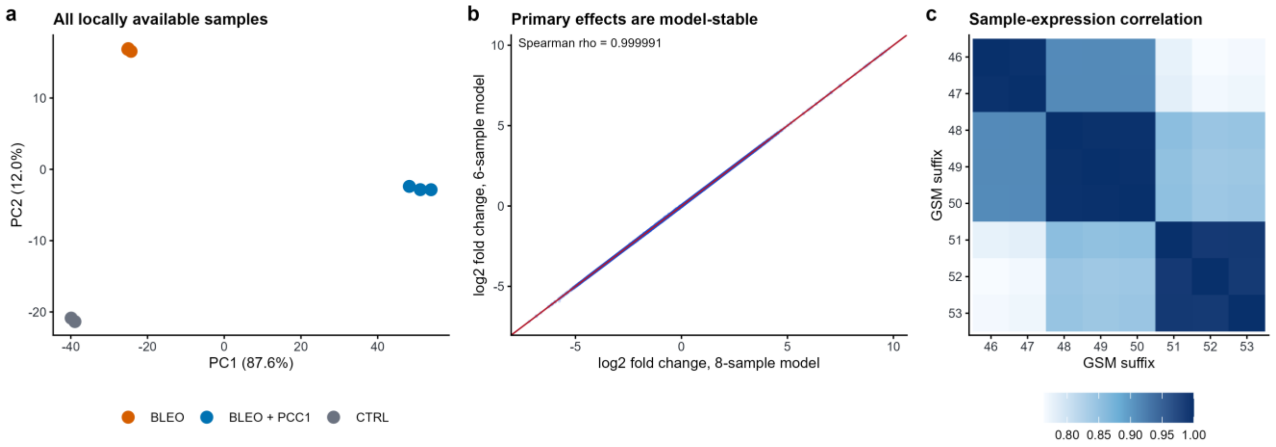


### Supplementary Figure S1 | Sample-level quality control and stability of the primary PCC1 contrast

**a,** PCA of all eight samples available in the NCBI combined count matrix: CTRL (n=2), BLEO (n=3) and BLEO+PCC1 (n=3). **b,** Concordance of log2 fold-change estimates from the complete six-sample primary model and the eight-sample sensitivity model containing the incomplete CTRL group. Spearman ρ=0.999991; the identity line is shown. **c,** Pairwise sample-correlation heatmap calculated from variance-stabilized expression. The expected third untreated-control sample, GSM4994845, was absent from the combined count matrix; all six samples required for the primary contrast were present.

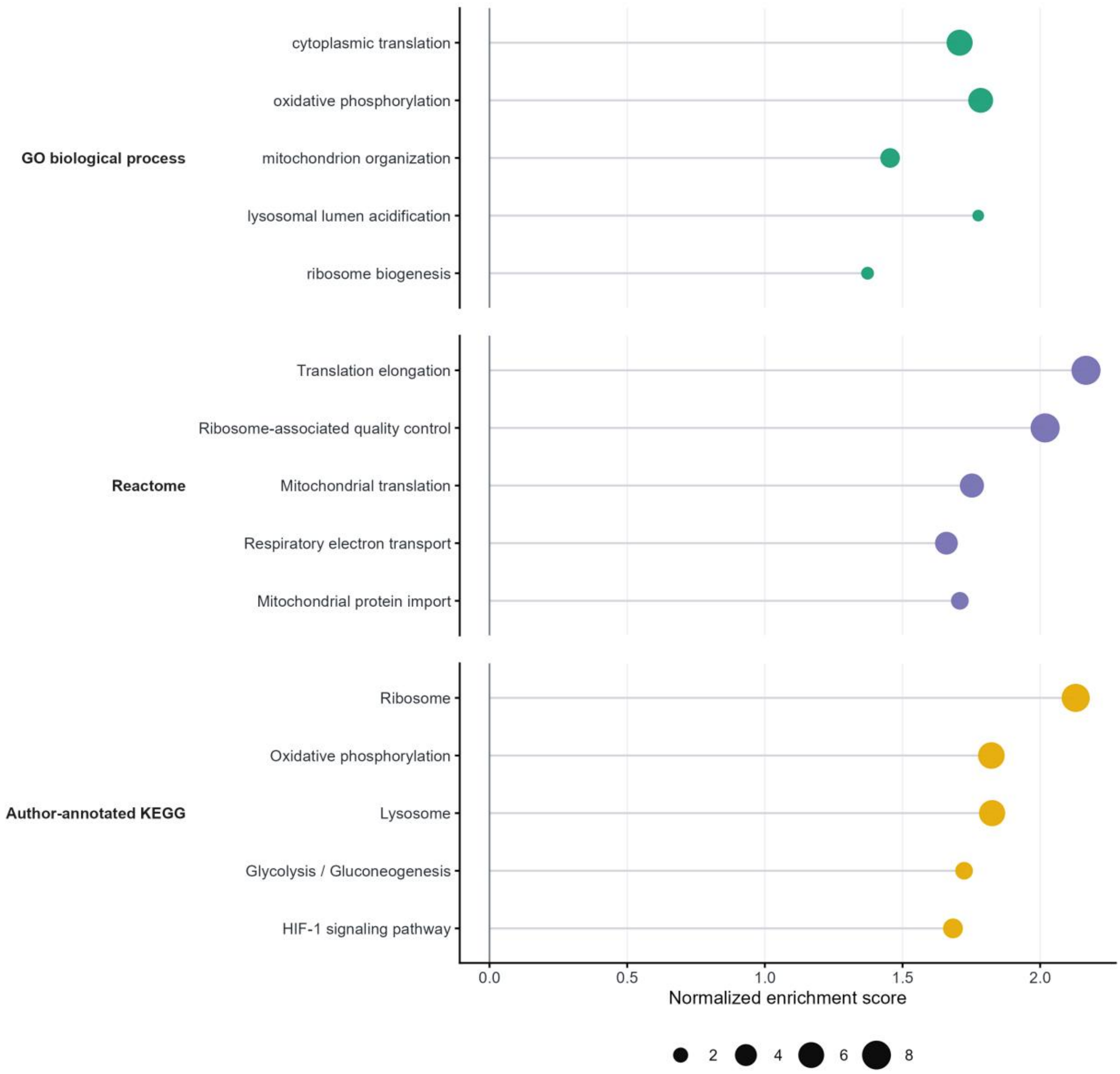


### Supplementary Figure S2 | Cross-annotation overview of cellular-infrastructure pathway enrichment

Representative positive ranked-enrichment results from GO Biological Process, Reactome and the KEGG annotations deposited with GSE164012. Position denotes NES and point size denotes -log10 FDR. Selected terms display non-redundant translation, ribosome-quality-control, mitochondrial, respiratory, lysosomal and metabolic themes. FDR values were calculated separately within each annotation collection.
